# Spatial cell graph analysis reveals skin tissue organization characteristic for cutaneous T cell lymphoma

**DOI:** 10.1101/2024.05.17.594629

**Authors:** Suryadipto Sarkar, Anna Möller, Anne Hartebrodt, Michael Erdmann, Christian Ostalecki, Andreas Baur, David B. Blumenthal

**Affiliations:** Biomedical Network Science Lab, Department Artificial Intelligence in Biomedical Engineering (AIBE), Friedrich-Alexander-Universität Erlangen-Nürnberg (FAU), Erlangen, Germany; Department of Dermatology, Uniklinikum Erlangen, Deutsches Zentrum Immuntherapie (DZI), Comprehensive Cancer Center Erlangen-European Metropolitan Area of Nuremberg (CCC ER-EMN), Friedrich-Alexander-Universität Erlangen-Nürnberg (FAU), Erlangen, Germany

## Abstract

Cutaneous T cell lymphomas (CTCLs) are non-Hodgkin lymphomas caused by malignant T cells which migrate to the skin. The cancerous T cells lead to rash-like lesions which can be difficult to distinguish from inflammatory skin conditions like atopic dermatitis (AD) and psoriasis (PSO). To characterize CTCL in comparison to these differential diagnoses, we carried out multi-antigen imaging on 69 skin tissue samples (21 CTCL, 23 AD, 25 PSO). The resulting spatially resolved protein abundance maps were then analyzed via scoring functions to quantify heterogeneity of the individual cells’ neighborhoods within spatial graphs inferred from the cells’ positions in the tissue samples (available as a Python package at https://github.com/bionetslab/SHouT). Our analyses reveal several characteristic patterns of skin tissue organization in CTCL, including a combination of increased local entropy and egophily as characteristic properties of spatial T cell neighborhoods in CTCL as compared to AD and PSO. These results could not only pave the way for high-precision diagnosis of CTCL, but may also facilitate further insights into cellular disease mechanisms.

## Introduction

Cutaneous T cell lymphomas (CTCL) present in the form of erythematous lesions, eruptions or patches on the skin. These lesions can both clinically and histologically resemble other non-cancerous inflammatory dermatological conditions, including atopic dermatitis (AD) and psoriasis (PSO)^1–4^, making diagnosis of CTCL challenging. Since CTCL needs to be treated thoroughly early on^5–9^, this is a major clinical problem. Techniques to reliably distinguish CTCL from the mimicking conditions AD and PSO therefore hold the promise to improve patient care in CTCL.

While several studies have explored the spatial heterogeneity of tumor microenvironment in CTCL and its potential relevance in prognosis^10,11^, a systematic quantification of spatial tissue heterogeneity in the context of CTCL does not exist. This prompted us to generate imaging-based spatially resolved protein abundance maps of skin tissue samples from CTCL, AD, and PSO patients treated at the University Hospital Erlangen, using multi-epitope ligand cartography (MELC)^12,13^. Based on a visual assessment of the images, tissue organisation was indeed altered in CTCL compared to AD and PSO samples. To quantify and objectivize this first subjective impression, we used the popular Squidpy package^14^ to generate graph representations of our data, where nodes are cells annotated with their cell type and edges encode spatial vicinity.

Since analysis with existing techniques available in Squidpy revealed only few differences, we developed a Python package called SHouT (short for “**s**patial **h**eter**o**geneity q**u**antification **t**ool”, available at https://github.com/bionetslab/SHouT), which allows to quantify tissue heterogeneity based on spatial cell graphs. Also beyond this study, SHouT can be used as an extension of the graph-based methods available in Squidpy to provide more nuanced insights into spatial tissue organization. SHouT revealed clear CTCL-specific characteristics of tissue organization, including higher mixing of cells of different types in the vicinity of T cells in CTCL as compared to AD and PSO. Randomization tests based on label permutation and subsampling verified the robustness of these findings.

## Results

### Overview of study design

Figure 1 provides an overview of our work (see Methods for details): Using MELC, we generated spatial protein abundance maps for 69 skin tissue samples (21 CTCL, 23 AD, 25 PSO). With this, we obtained images of resolution 2018 *×* 2018 pixels for at least 35 protein channels per sample, leading to over 140 million pixel values. We then carried out cell segmentation for all images, using the propidium iodide and CD45 channels as markers for nucleus and cell membrane, respectively. Subsequently, adaptive thresholding was used to quantify protein abundances within the individual cells, and cell types were assigned via a rule-based marker gene approach (Supplementary Figure 1), using skin tissue single-cell RNA-sequencing (scRNA-seq) data from the Human Protein Atlas (HPA)^15^ as reference.

**Figure 1.**
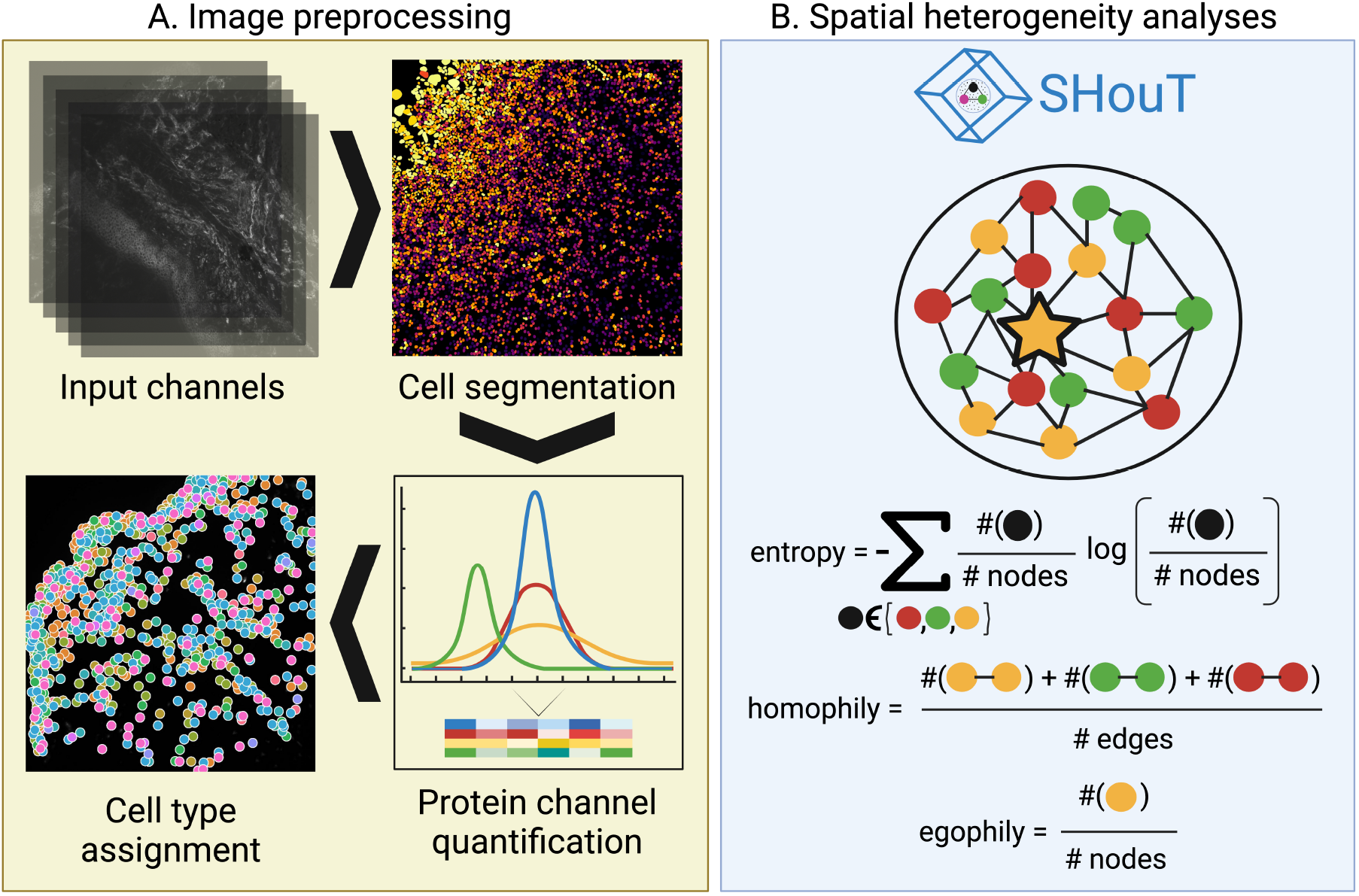
Overview of our analyses. (A) We generated multi-antigen images for skin tissue samples from 21 CTCL, 23 AD, and 25 PSO patients. Subsequently, images were pre-processed via cell segmentation, cell-level protein abundance quantification, and cell type assignment. (B) We then computed spatial graph representations for all samples, which we analyzed using SquidPy as well as different heterogeneity scores implemented in our Python package SHouT: local and global entropy, local and global homophily, and egophily.

Next, we computed a spatial graph for each sample, where two cells are connected by an edge if they are adjacent to each other in the spatial map. The spatial graphs were then analyzed with SquidPy’s `centrality_scores` and `nhood_enrichment` functions, which compute degree and closeness centralities of the individual cells within the spatial graphs and enrichment of cell type co-occurrence within one-hop spatial neighborhoods. Moreover, we analyzed the cell graphs with our novel Python package SHouT. SHouT provides two sample-level scores that quantify overall tissue heterogeneity:

- The edge-based *global homophily* score represents the fraction of edges in the spatial graph that connect cells of the same type. A high score indicates low overall heterogeneity.
- The node-based *global entropy* score represents how well-balanced the numbers of cells of each cell type are throughout the network. A high score indicates high overall heterogeneity.

Moreover, SHouT provides three cell-level scores that quantify tissue heterogeneity within the *r*-hop neighborhood of an individual cell *c* in the spatial graphs (Figure 1B):

- The edge-based *local homophily* score represents the fraction of edges in the *r*-hop neighborhood of *c* that connect cells of the same type. A high score indicates low heterogeneity in the tissue region surrounding cell *c*.
- The node-based *local entropy* score represents how well-balanced the numbers of cells of each cell type are in the *r*-hop neighborhood of *c*. A high score indicates high heterogeneity in the tissue region surrounding cell *c*.
- The node-based *egophily* score represents the fraction of cells within the *r*-hop neighborhood of *c* that have the same cell type as *c*. A high score indicates low heterogeneity in the tissue region surrounding cell *c*.

By specifying the radius *r*, the user can select the desired granularity of the cell-level scores. With *r* = 1, only cells in the immediate vicinity of *c* are considered. On the other extreme, setting *r* to the diameter of the spatial graph renders local homophily and entropy equivalent to global homophily and entropy, respectively. Setting *r* to intermediate values allows quantification of meso-scale patterns in the spatial graphs.

Figure 2 provides a high-level overview of the dataset. Figure 2A shows the average cell type composition stratified by sample type (see Supplementary Figure 2 for underlying distributions of sample-specific cell type fractions). With the exception of increased abundance of endothelial cells in AD samples and increased abundance of macrophages in PSO samples, we did not observe strong differences in cell type composition between the three conditions, emphasizing the need for more in-depth analyses. Figure 2B shows the propidium iodide channel used for cell nucleus identification for one sample. Figure 2C shows the propidium iodide channel for the same sample, with the CD4 channel overlayed on top, as an example for the data used to quantify protein abundances.

**Figure 2.**
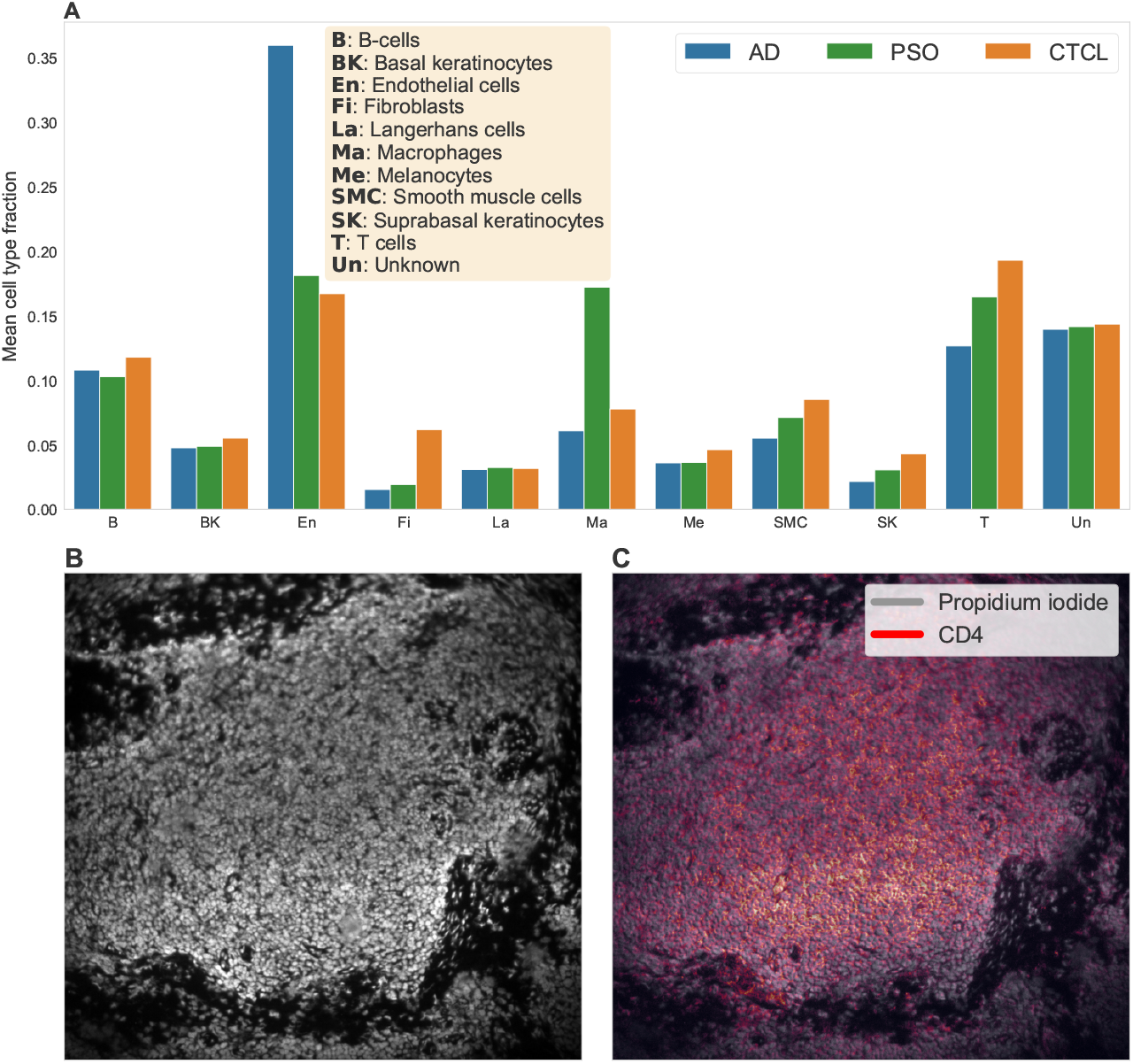
Summary of the multi-antigen imaging dataset used for this study. (A) Mean cell type fractions per condition, across all of the samples. (B) The propidium iodide channel used for cell segmentation for one sample. (C) CD4 expression overlayed on the propidium iodide channel.

### Heterogeneity analysis reveals CTCL-specific patterns in the neighborhoods of T cells and basal keratinocytes

Figure 3 shows the results of analyzing the spatial graphs with SHouT and SquidPy’s functions `centrality_scores` and `nhood_enrichment`. All shown *P*-values were computed with the two-sided Mann-Whitney U (MWU) test and were Bonferroni-corrected with respect to the number of tests per score type (neighborhood enrichment scores: number of condition pairs *×* number of cell type pairs; centrality scores and global heterogeneity scores: number of condition pairs; local heterogeneity scores: number of condition pairs *×* number of cell types *×* number of tested radii (5)).

**Figure 3.**
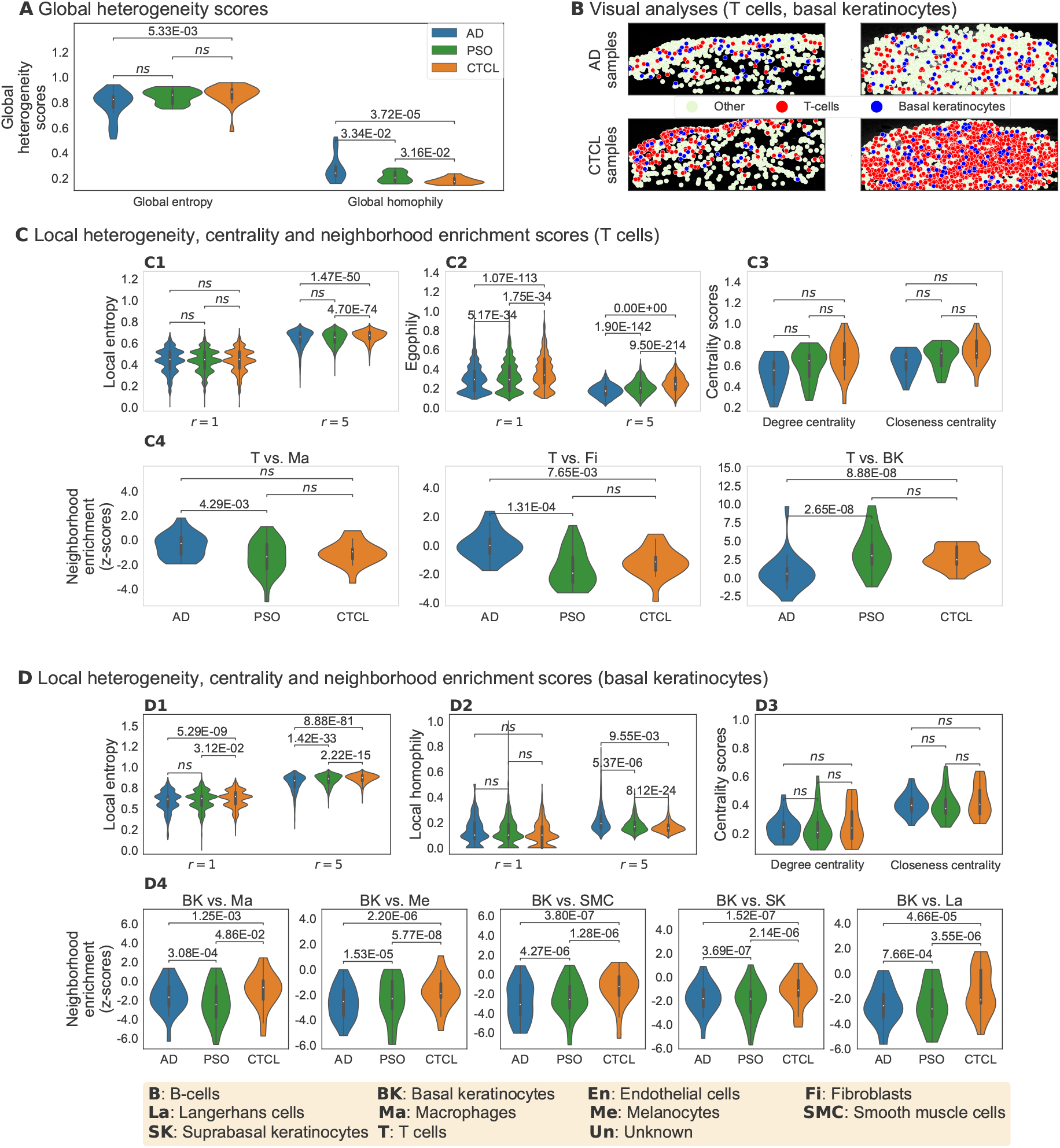
Results of spatial analyses of AD, PSO, and CTCL samples with adjusted MWU *P*-values. The most important differences between CTCL samples as compared to AD or PSO are elevated local entropy and egophily scores for neighborhoods of T cells and elevated local entropy and local homophily scores for neighborhoods of basal keratinocytes. (A) Global heterogeneity scores. (B) Visualization of spatial distributions of T cells and basal keratinocytes in two AD and two CTCL samples. (C) Selected local heterogeneity, centrality, and neighborhood enrichment scores for T cells. (D) Selected local heterogeneity, centrality, and neighborhood enrichment scores for basal keratinocytes.

The two global heterogeneity scores show that overall tissue heterogeneity is highest in CTCL (lowest global homophily, largest global entropy), followed by PSO and AD (Figure 3A). When focusing on T cells (Figure 3C), we observe significantly elevated local entropy (Figure 3C1) and egophily (Figure 3C2) scores in CTCL. That is, in skin samples from CTCL patients, T cells tend to cluster together (unsurprisingly, since the tumour cells are T cells) and at the same time are surrounded by tissue that exhibits a higher mixing of cell types than the spatial neighborhoods of T cells in AD or PSO (increased local entropy). Interestingly, the differences are more pronounced and the score distributions are smoother for radius *r* = 5 than for *r* = 1, highlighting the importance of incorporating *r*-hop neighborhoods into SHouT’s local heterogeneity scores (see Supplementary Figures 3, 4 and 5 for additional results with *r* = 5 for all cell types and Supplementary Figure 6 for distributions of ShouT scores across radii *r* ∈ {1, 2, 3, 4, 5}).

In contrast to SHouT, Squidpy’s `centrality_scores` function did not reveal significant differences between the three conditions (Figure 3C3, see Supplementary Figure 7 for additional results corresponding to all cell types). Neighborhood enrichment analysis with Squidpy’s `nhood_enrichment` function (Figure 3C4) did reveal significant differences in co-occurrence between T cells and, respectively, macrophages, fibroblasts, and basal keratinocytes. However, the observed differences are much smaller than for local entropy and homophily and no clear picture emerges that would allow to robustly distinguish CTCL samples from AD and PSO samples based on neighborhood enrichment scores of T cells.

We obtained highly significant differences in SHouT’s local heterogeneity scores also for another cell type besides T cells, namely, basal keratinocytes (Figure 3D). Like for T cells, local entropy is increased in CTCL in comparison to AD and PSO (Figure 3D1), i. e., tissue in the vicinity of basal keratinocytes exhibits a higher cell type mixing in CTCL than in the two other conditions. Moreover, and in contrast to the results for T cells, we simultaneously observe elevated local homophily scores for basal keratinocytes (Figure 3D2). That is, tissue in the vicinity of basal keratinocytes exhibits a higher co-localization of cells of the same type in CTCL than in PSO or AD (increased local homophily), even though the cell type heterogeneity is increased (increased local entropy). This shows that the seemingly similar local entropy and local homophily scores indeed quantify distinct properties of spatial tissue organization. While Squidpy’s `centrality_scores` function did not reveal significant differences related to basal keratinocytes (Figure 3D3), neighborhood enrichment analysis showed that, in CTCL samples, occurrence of macrophages, melanocytes, smooth muscle cells, suprabasal keratinocytes, and Langerhans cells is increased in the vicinity of basal keratinocytes (Figure 3D4).

### The identified CTCL-specific patterns are robust to label permutation and subsampling

We carried out permutation tests to assess the reliability of the identified differences in the local heterogeneity scores with radius *r* = 5 shown in Figure 3. Specifically, we shuffled the condition labels across all 69 samples and then used the MWU test to compare the local heterogeneity scores between the obtained randomized conditions. We repeated this for 100 iterations, leading to 100 *P*-values per cell type and condition pair. Figure 4 shows the resulting *P*-value distributions (histograms), together with the *P*-values for the original condition labels (red lines). For all eleven combinations of scores and condition pairs where results are significant for the original, unshuffled condition labels (all plots in Figure 4 except for the one at the bottom left), the *P*-values obtained for the original condition labels are much smaller than those obtained for the shuffled labels. This indicates that the heterogeneity scores indeed reveal robust differences between CTCL, AD, and PSO.

**Figure 4.**
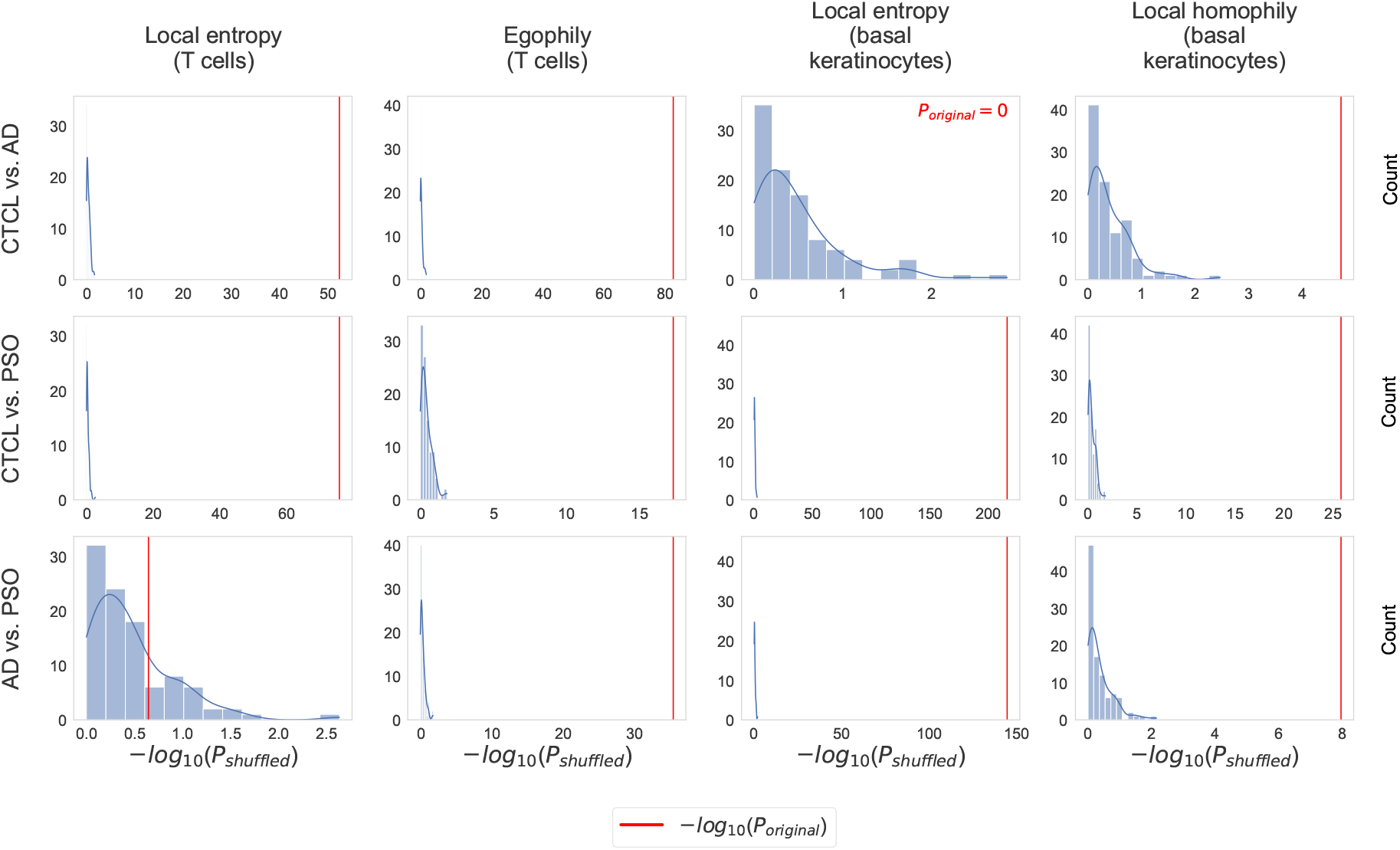
Results of permutation tests to assess the robustness of the differences in the local heterogeneity scores with radius *r* = 5 shown in Figure 3. All *P*-values were computed with the MWU test. Unlike for the *P*-values shown in Figure 3, we did not apply Bonferroni correction to adjust for multiple testing (since we are comparing *P*-values, adjusting for multiple testing leaves the results invariant).

Moreover, we carried out subsampling to assess if the identified CTCL-specific tissue organization patterns are robust to varying compositions of the CTCL, AD, and PSO cohorts. Specifically, we subsampled 15 samples for each of the three conditions. Subsequently, for each condition pair, we computed MWU *P*-values based on the SHouT’s local heterogeneity scores (with *r* = 5) for T cells and basal keratinocytes, using only the samples from the subsampled patients. We repeated this 100 times, leading to 100 MWU *P*-values for each condition pair, considered cell type, and SHouT score. The resulting *P*-value distributions (boxplots) are shown in Figure 5, together with the *P*-values obtained when using all samples (red lines) and the Bonferonni-corrected significance cutoff *P*_cutoff_ = 0.05*/*(number of condition pairs *×* number of cell types *×* number of tested radii) (blue lines). For all cases where the original *P*-values are significant (red lines to the right of blue lines), all or the vast majority of the *P*-values obtained upon subsampling are significant, too. This shows that SHouT reveals CTCL-specific tissue patterns that are robust to subsampling.

**Figure 5.**
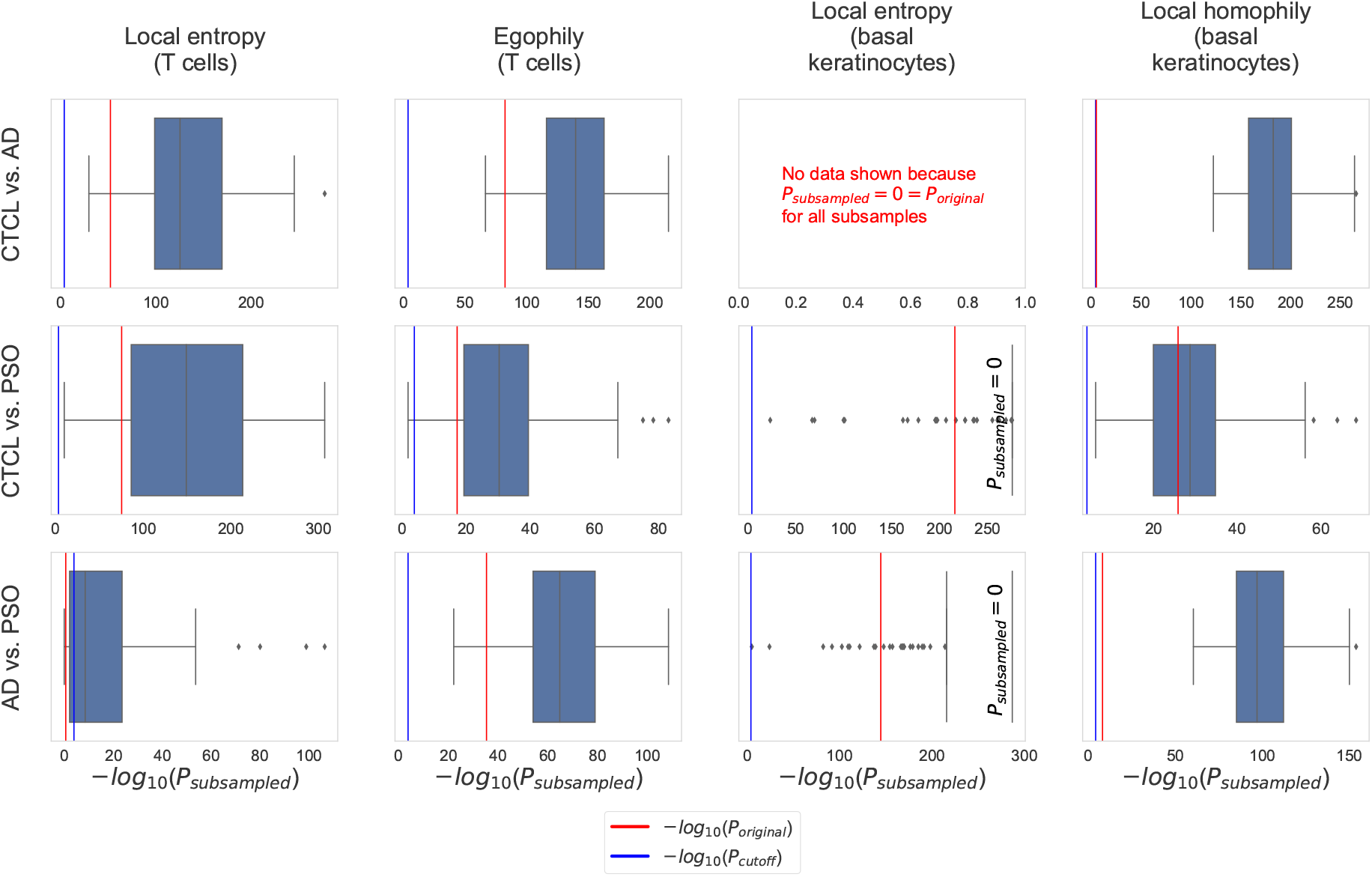
Results of subsampling tests to assess the robustness of the differences in the local heterogeneity scores with radius *r* = 5 shown in Figure 3. All *P*-values were computed with the MWU test. The significance cutoff is Bonferroni-adjusted, i. e., *P*_cutoff_ = 0.05*/*(number of condition pairs *×* number of cell types *×* number of tested radii) = 0.05*/*(3 *×* 11 *×* 5). *P*-values are non-adjusted.

### SHouT scales to samples with large numbers of cells

To ensure usability, runtime efficiency is an important property of data-centric methods for the analysis of biomedical data. We therefore systematically tested SHouT with respect to its runtime requirements when run on samples with varying numbers of cells or when run with varying radii. The results are shown in Figure 6. We observe that SHouT scales linearly with the numbers of cells per sample, achieving runtimes of little more than a minute even for the samples with the highest cell counts in our dataset (Figure 6A). Increasing the radius *r* only marginally increases the runtime, showing that SHouT’s heterogeneity scores can be computed efficiently independently of the choice of *r* (Figure 6B).

**Figure 6.**
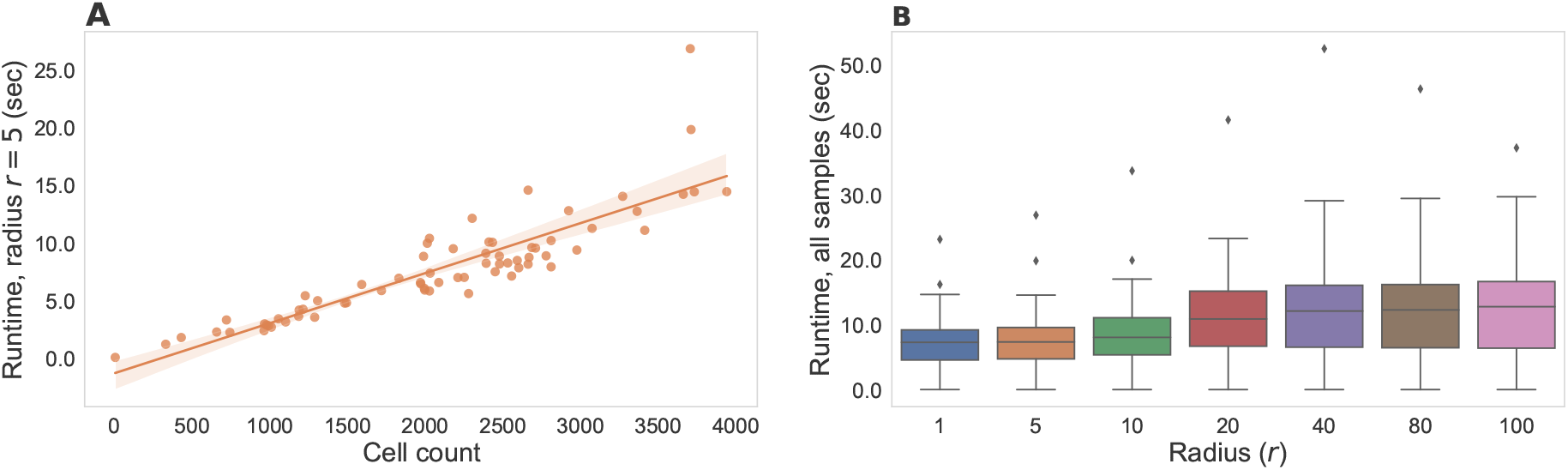
Results of scalability tests for our Python package SHouT. Runtime measurements include all subroutines detailed in the “Quantification of spatial tissue heterogeneity with SHouT” subsection of the Methods: construction of the spatial neighborhood graph *G* = (*V, E, λ*_*V*_), computation of the two global scores *H*(*G*) and *h*(*G*), and computation of the three local scores *H*_*r*_(*c*), *h*_*r*_(*c*), and *e*_*r*_(*c*) for all cells *c ∈ V* and a fixed radius *r*. To ensure a stable execution environment, tests were run on a Linux compute server with 500 GB of main memory and four AMD EPYC 7402 24-core processors with 1.5 Ghz (without using multi-threading). However, SHouT does not require such large-scale ressources and can be run on a standard laptop. (A) Runtimes with a fixed radius *r* = 5 for all samples in our dataset. (B) Runtime distributions for varying radii across all samples in our dataset.

## Discussion

Our analyses identified CTCL-specific patterns of tissue organisation as compared to PSO and AD in the vicinity of T cells and basal keratinocytes. Since CTCL is a T cell malignancy, observing characteristic patterns in the vicinity of T cells is not too surprising. In fact, existing studies suggest that malignant T cells and their cross-talk with other cells induce disorganization in the epidermal architecture^16–18^, which is well aligned with our findings. Also our results for basal keratinocytes are plausible in the light of the literature. For instance, several studies have identified hyperproliferation and/or de-differentiation of keratinocytes in CTCL^17,19,20^.

An important aspect of our findings is that the SHouT scores underlying our results are (1) purely quantitative, (2) interpretable by design (each SHouT score has a natural interpretation that can be explained with few sentences), and (3) deterministically computed (computing the SHouT scores does not involve randomized subroutines). Their purely quantitative nature sets our SHouT scores apart from the current standard of care in CTCL diagnosis, which is typically based on (subjective) expert opinions. Their interpretability is a decisive advantage over predictions provided by deep learning models, which are often perceived as black boxes^21^. And the fact that SHouT is deterministic ensures that it consistently computes the same results when run several times on equivalent input — unlike many other methods in data-centric biomedicine^22,23^ including Squidpy’s `nhood_enrichment` function. Together, these three properties make the SHouT scores ideal ingredients for potential future biomarkers based on spatial omics data, both in the CTCL use case presented here and beyond.

However, several limitations of our study have to be addressed before translation into clinical care may become feasible. First and foremost, the MELC technology used to generate the data underlying this study is highly non-standardized (only few prototypes exist worldwide, two of them at UKER). This had strong implications for the data analysis pipeline used for this study, where we had to resort to customized solutions for almost all pre-processing steps (cell segmentation, protein abundance quantification, cell type assignment). While we are confident that the methodological choices we made for individual steps in our pipeline are adequate for the data, they still introduce a certain amount of contingency which may distort the results of our analyses in a hard-to-control way. Before translation into clinical care can become an option, it would hence be important to see if our results can be reproduced for spatial omics generated with more established technologies (e. g., MERFISH^24^ or MIBI^25^), for which validated pre-processing pipelines are available.

## Methods

### Sample collection and multi-antigen imaging

69 skin tissue samples (21 CTCL, 23 AD, 25 PSO) from a total of 27 treated patients (8 CTCL, 7 AD, 12 PSO) were collected at the University Hospital Erlangen. The study has been approved by the Ethics Committee of the Medical Faculty of the Friedrich-Alexander-Universität Erlangen-Nürnberg (approval date: July 5, 2023; approval number: 23-132-B). We used the MELC technology^12,13,26^ to generate multi-antigen imaging data. MELC efficiently combines seamless assimilation of molecular and anatomical information *in situ* by employing a cyclic process of three steps: (1) protein-specific fluorescent antigen staining, (2) imaging, and (3) photobleaching. The workflow is completely automated, performing multi-antigen imaging on a fixed and mounted tissue or cell sample, without the requirement of any human involvement. Thanks to this automated workflow, post imaging, the individual protein channels can be mapped into one consolidated tissue map whilst preserving spatial information. In order to assemble a set of antibodies with a strong and specific staining pattern, we screened over 500 antibodies on CTCL tissue. This unbiased approach yielded 36 antibodies (Supplementary Table 1).

### Cell segmentation and protein abundance quantification

Owing to the properties of DNA- and histone-binding, propidium iodide is pervasively used to stain cell nuclei in fluorescent microscopy^27^. We therefore used the propidium iodide channel for cell nucleus segmentation, relying on a pre-trained model provided by the `stardist.models.StarDist2D` function of the popular cell detection library StarDist^28,29^.

For cell membrane segmentation, we used the channel for the transmembrane protein tyrosine phosphate (CD45), which is not only present in all nucleated hematopoietic cells but is also one of the most commonly found membrane proteins in such cells^30^. However, owing to the often non-convex shape of cell membranes as opposed to cell nuclei in our dataset, the StarDist model under-segments the cytoplasm. As a workaround, for all nuclei where the StarDist model managed to automatically identify the cell membrane, we calculated an average ratio between radii of cells and nuclei. We then used this average ratio to draw a circle around each of the segmented nuclei for which the StarDist model did not identify a cell membrane. If two such circles overlapped, pixels within the intersection where assigned to the closest nucleus, thereby ensuring that all cell segments are pairwise disjoint.

Transitioning from pixel intensity to cell-level protein abundances was challenging due to the following factors: (1) MELC images are especially susceptible to salt-and-pepper noise. (2) Different protein channels have different intensity levels. (3) Even within the same channel, different regions often have different levels of intensity. Because of these factors, there is no single mapping which would allow to uniformly compute cell-level protein abundance scores based on pixel intensities across all samples and sample regions. We therefore made use of adaptive thresholding which helps circumvent this issue by breaking the image into smaller windows comprising fewer pixels, subsequently computing a different threshold for every window. Specifically, we used OpenCV’s (https://opencv.org/) `cv2.adaptiveThreshold` function to perform adaptive thresholding. This function binarizes the individual channels by setting each pixel to 1 if its intensity is above the Gaussian weighted mean of its vicinity (here, a window of size 201 *×* 201) plus a constant *C* (here, the standard deviation of the intensity values across the current protein channel). We then transformed the binarized images into floating point protein abundance matrices *A* by setting the abundance *A*(*c, p*) of protein *p* in cell *c* to the fraction of positive binarized pixels within the segment corresponding to *c*.

### Cell type assignment

Since only few exemplars of the MELC imaging system exist, automated cell type annotation tools for our data do not exist. Moreover, initial tests showed that cell type annotation tools developed for data generated by other multiplexing platforms are not applicable for our data. We therefore developed a simple rule-based cell type annotation workflow, making use of single-cell RNA sequencing (scRNA-seq) data specific to skin tissue from HPA as reference. Specifically, we downloaded normalized gene expression values *X* (*C, g*), where *C* denotes cell clusters provided in HPA and *g* denotes genes. Moreover, we made use of HPA’s cell type annotations *σ* (*C*) *∈* Σ, where Σ is the set of cell types for skin tissue used by HPA. Since, in HPA, there are cell types to which several clusters are assigned, we computed cell type-specific gene expression values

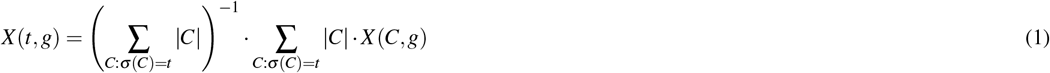

via weighted mean aggregation, for all pairs of cell types *t* and genes *g*.

Let 𝒞 be the set of all cells contained in any of the samples for any of the three conditions CTCL, PSO, and AD, 𝒢 be the set of all genes encoding the measured proteins, and *A*(*c, g*) be the abundance of the protein encoded by *g* in the cell *c*. Our cell type assignment workflow iteratively assigns cell types *t*^*⋆*^ *∈* Σ to a subset 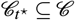 of the cells. Upon assigning the cell type *t*^*⋆*^ to the cells contained in 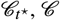 and Σ are updated as 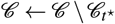 and Σ *←* Σ \ *{t*^*⋆*^*}*, respectively. The process stops when either 𝒞 = Ø, Σ = Ø, or there are no good genes (defined below). In the latter two cases, the remaining cells in 𝒞 are assigned the cell type label “unknown”.

To find *t*^*⋆*^ and 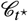 within one iteration, we computed HPA-based spread scores

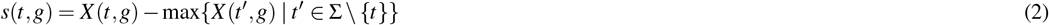

for all pairs of not yet assigned cell types *t ∈* Σ and genes *g ∈* 𝒢 and sorted the pairs in decreasing order of *s*(*t, g*), leading to a sorted list *L*. Genes *g* with large spread scores *s*(*t, g*) are potential marker genes for the cell type *t*, based on the scRNA-seq data in HPA. Next, we fit a bimodal Gaussian mixture models to the vectors of protein abundances (*A*(*c, g*))_*c∈𝒞*_ for the not yet assigned cells (using scikit-learns’s `GaussianMixture` class) and all genes *g ∈* 𝒢 and assessed if the obtained distributions are indeed bimodal. For this, we checked the condition

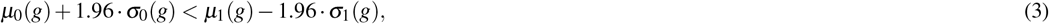

where *μ*_0_(*g*) and *σ*_0_(*g*) are the mean and standard deviation values for the mode with the lower mean and *μ*_1_(*g*) and *σ*_1_(*g*) are the mean and standard deviation values for the mode with the higher mean. Genes for which the condition holds are called “good”. When we found at least one good gene, we picked the first pair (*t*^*⋆*^, *g*^*⋆*^) from the sorted list *L* for which *g*^*⋆*^ is good. Then, we assigned the cell type *t*^*⋆*^ to the cells 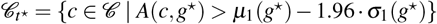 and continued with the next iteration of our cell type assignment protocol. When no good gene was found (for our data, this happened when “granulocytes” was the only cell type label left in Σ), we stopped the protocol.

The resulting rule-based cell type assignment tree is shown in Supplementary Figure 1. Heatmaps showing the corresponding (A) cell type-averaged gene expression values from HPA and (B) protein abundances from our data, are shown in Supplementary Figure 8. In the following, we let *T* denote the set of used cell type labels (in our case: the label “unknown” and all labels contained in Σ except “granulocytes”) and let *λ* : 𝒞 *→ T* denote the constructed cell type label function.

### Quantification of spatial tissue heterogeneity with SHouT

SHouT starts by computing sample-specific spatial neighborhood graphs *G* = (*V, E, λ*_*V*_) from the pre-processed imaging data, where *V ⊆* 𝒞 is the set of cells for the sample under consideration, the set *E* contains an edge *cc*^*′*^ for two cells *c, c*^*′*^ *∈ V* if *c* and *c*^*′*^ are spatially adjacent (computed with Squidpy’s `spatial_neighbors` function with the parameter `delaunay` set to `True`), and *λ*_*V*_ denotes the restriction of the cell type label function *λ* to *V*. Based on the spatial graphs, SHouT computes two global scores that quantify heterogeneity for the entire graph *G* and three local, cell-specific scores.

The first global score — global (normalized) entropy — is defined as

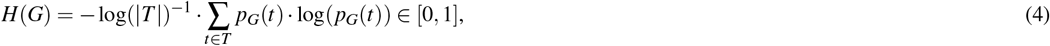

where *p*_*G*_(*t*) = |*{c ∈ V* | *λ*_*V*_ (*c*) = *t}*|*/*|*V* | is the fraction of cells in *V* that are of type *t*. Large values of *H*(*G*) indicate that cell type heterogeneity is high for the sample represented by *G*. The second global score — global homophily — is defined as the fraction of edges

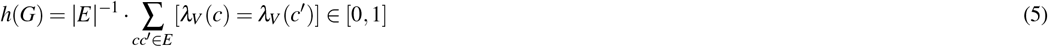

in the spatial graph *G* that connect cells of the same type ([*·*] : {True, False} → {0, 1*}* is the Iverson bracket, i. e., [True] = 1 and [False] = 0). Large values of *h*(*G*) indicate that cells tend to be adjacent to cells of the same type in the sample represented by *G*.

In addition to the two global scores, SHouT provides three local scores to quantify heterogeneity within the *r*-hop neighborhood *N*_*r*_(*c*) = *{c*^*′*^ *∈ V* | *d*_*G*_(*c, c*^*′*^ *≤ r}* of an individual cell *c ∈ V*. Here, *r* is a hyper-parameter and *d*_*G*_ : *V ×V →* ℕ is the shortest path distance. The first local score — local (normalized) entropy — is defined as follows:

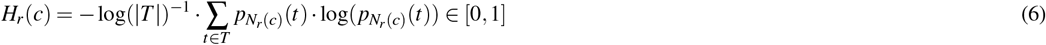

The only difference to its global counterpart is that the cell type fractions 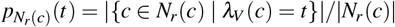 are computed only with respect to the cells contained in the *r*-hop neighborhood of *c*. Also the second local score — local homophily — is defined in a similar way as the global version:

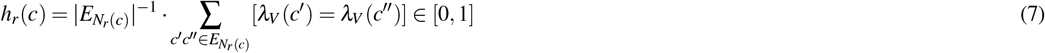

Here, the difference to the global version is that we only consider the subset of edges 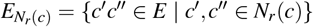 that connect two cells contained in the *r*-hop neighborhood of *c*. The last local score — egophily — does not have a global counterpart. It is defined as the fraction

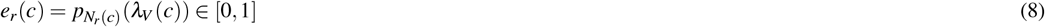

of cells within the *r*-hop neighborhood of *c* that have the same cell type as *c*. SHouT provides very efficient vectorized implementations of all heterogeneity scores, relying on SciPy’s `sparse.csgraph.shortest_path` function for fast computation of shortest path distances.

## Data availability

The MELC data underlying this study are available on Zenodo: https://doi.org/10.5281/zenodo.11125482. Pre-clustered, skin-specific scRNA-seq data used as reference for cell type assignment are available at https://www.proteinatlas.org/download/rna_single_cell_type_tissue.tsv.zip. The corresponding cluster annotations can be obtained at https://www.proteinatlas.org/download/rna_single_cell_cluster_description.tsv.zip.

## Code availability

The code of the SHouT Python package is available at https://github.com/bionetslab/SHouT. Code to reproduce the results reported in this paper is available at https://github.com/bionetslab/ctcl_case_study.

## Acknowledgements

D.B.B. and A.M. were funded by the Deutsche Forschungsgemeinschaft (DFG, German Research Foundation) – 516188180. Figure 1 and Supplementary Figure 1 were generated with BioRender.com. D.B.B. and A.H. were supported by the German Federal Ministry of Education and Research (BMBF, grant no. 031L0309A).

## Author contributions statement

S.S. and D.B.B. conceived and designed this study, implemented SHouT, analysed the results, and drafted the manuscript. S.S. carried out the analyses. C.O. generated the data. C.O. and M.E. annotated the data. A.M. pre-processed the data. D.B.B., A.B., and A.H. supervised this work. All authors reviewed and approved the manuscript.

## References

1. Semaan, S., Abel, M. K., Raffi, J. & Murase, J. E. A clinician’s guide to cutaneous T-cell lymphoma presenting as recalcitrant eczematous dermatitis in adults. Int. J. Womens. Dermatol. 7, 422–427, DOI: 10.1016/j.ijwd.2021.04.004 (2021).

2. Tomasini, C., Novelli, M., Fanoni, D. & Berti, E. F. Erythema multiforme-like lesions in primary cutaneous aggressive cytotoxic epidermotropic CD8+ T-cell lymphoma: A diagnostic and therapeutic challenge. J. Cutan. Pathol. 44, 867–873, DOI: 10.1111/cup.12995 (2017).

3. Papadaki, M., Saraki, K., Karagianni, F., Piperi, C. & Papadavid, E. Cutaneous T-cell lymphoma: aetiopathogenesis and current diagnostic and therapeutic developments. Eur. J. Dermatol. DOI: 10.1684/ejd.2020.3712 (2020).

4. Roediger, B. & Schlapbach, C. T cells in the skin: Lymphoma and inflammatory skin disease. J. Allergy Clin. Immunol. 149, 1172–1184, DOI: 10.1016/j.jaci.2022.02.015 (2022).

5. Willemze, R. et al. The 2018 update of the WHO-EORTC classification for primary cutaneous lymphomas. Blood 133, 1703–1714, DOI: 10.1182/blood-2018-11-881268 (2019).

6. Hristov, A. C., Tejasvi, T. & A Wilcox, R. Cutaneous T-cell lymphomas: 2021 update on diagnosis, risk-stratification, and management. Am. J. Hematol. 96, 1313–1328, DOI: 10.1002/ajh.26299 (2021).

7. Franceschi, J. et al. Survival and prognostic factors in patients with aggressive cutaneous T-cell lymphomas. Acta Derm. Venereol. 102, adv00676, DOI: 10.2340/actadv.v102.1087 (2022).

8. Brunner, P. M., Jonak, C. & Knobler, R. Recent advances in understanding and managing cutaneous T-cell lymphomas. F1000Res. 9, DOI: 10.12688/f1000research.21922.1 (2020).

9. Miyagaki, T. Diagnosis and prognostic stratification of cutaneous lymphoma. J. Dermatol. 49, 210–222, DOI: 10.1111/1346-8138.16099 (2022).

10. Kalliara, E., Belfrage, E., Gullberg, U., Drott, K. & Ek, S. Spatially guided and single cell tools to map the microenvironment in cutaneous T-Cell lymphoma. Cancers 15, DOI: 10.3390/cancers15082362 (2023).

11. Feng, Y. et al. Spatial transcriptomics reveals heterogeneity of macrophages in the tumor microenvironment of granulomatous slack skin. J. Pathol. 261, 105–119, DOI: 10.1002/path.6151 (2023).

12. Schubert, W. et al. Analyzing proteome topology and function by automated multidimensional fluorescence microscopy. Nat. Biotechnol. 24, 1270–1278, DOI: 10.1038/nbt1250 (2006).

13. Schubert, W. Topological proteomics, toponomics, MELK-technology. Adv. Biochem. Eng. Biotechnol. 83, 189–209, DOI: 10.1007/3-540-36459-5_8 (2003).

14. Palla, G. et al. Squidpy: a scalable framework for spatial omics analysis. Nat. Methods 19, 171–178, DOI: 10.1038/s41592-021-01358-2 (2022).

15. Karlsson, M. et al. A single-cell type transcriptomics map of human tissues. Sci Adv 7, DOI: 10.1126/sciadv.abh2169 (2021).

16. Thumann, P. et al. Interaction of cutaneous lymphoma cells with reactive T cells and dendritic cells: implications for dendritic cell-based immunotherapy. Br. J. Dermatol. 149, 1128–1142, DOI: 10.1111/j.1365-2133.2003.05674.x (2003).

17. Thode, C. et al. Malignant T cells secrete galectins and induce epidermal hyperproliferation and disorganized stratification in a skin model of cutaneous t-cell lymphoma. J. Invest. Dermatol. 135, 238–246, DOI: 10.1038/jid.2014.284 (2015).

18. Stolearenco, V. et al. Cellular interactions and inflammation in the pathogenesis of cutaneous T-Cell lymphoma. Front. Cell. Dev. Biol. 8, 851, DOI: 10.3389/fcell.2020.00851 (2020).

19. Weiss, R. A., Eichner, R. & Sun, T. T. Monoclonal antibody analysis of keratin expression in epidermal diseases: a 48- and 56-kdalton keratin as molecular markers for hyperproliferative keratinocytes. J. Cell Biol. 98, 1397–1406, DOI: 10.1083/jcb.98.4.1397 (1984).

20. Edelson, R. L. Cutaneous T cell lymphoma: the helping hand of dendritic cells. Ann. N. Y. Acad. Sci. 941, 1–11, DOI: 10.1111/j.1749-6632.2001.tb03705.x (2001).

21. Qamar, T. & Bawany, N. Z. Understanding the black-box: towards interpretable and reliable deep learning models. PeerJ Comput. Sci 9, e1629, DOI: 10.7717/peerj-cs.1629 (2023).

22. Bernett, J. et al. Robust disease module mining via enumeration of diverse prize-collecting steiner trees. Bioinformatics 38, 1600–1606, DOI: 10.1093/bioinformatics/btab876 (2022).

23. Sarkar, S. et al. Online bias-aware disease module mining with ROBUST-Web. Bioinformatics 39, btad345, DOI: 10.1093/bioinformatics/btad345 (2023).

24. Chen, K. H., Boettiger, A. N., Moffitt, J. R., Wang, S. & Zhuang, X. RNA imaging. spatially resolved, highly multiplexed RNA profiling in single cells. Science 348, aaa6090, DOI: 10.1126/science.aaa6090 (2015).

25. Ptacek, J. et al. Multiplexed ion beam imaging (MIBI) for characterization of the tumor microenvironment across tumor types. Lab. Invest. 100, 1111–1123, DOI: 10.1038/s41374-020-0417-4 (2020).

26. Schubert, W. Exploring molecular networks directly in the cell. Cytom. A 69, 109–112, DOI: 10.1002/cyto.a.20234 (2006).

27. Martin, R. M., Leonhardt, H. & Cardoso, M. C. DNA labeling in living cells. Cytom. A 67, 45–52, DOI: 10.1002/cyto.a.20172 (2005).

28. Schmidt, U., Weigert, M., Broaddus, C. & Myers, G. Cell detection with Star-Convex polygons. In Medical Image Computing and Computer Assisted Intervention – MICCAI 2018, 265–273, DOI: 10.1007/978-3-030-00934-2_30 (Springer International Publishing, 2018).

29. Weigert, M., Schmidt, U., Haase, R., Sugawara, K. & Myers, G. Star-convex polyhedra for 3D object detection and segmentation in microscopy. In 2020 IEEE Winter Conference on Applications of Computer Vision (WACV), vol. 0, 3655–3662, DOI: 10.1109/WACV45572.2020.9093435 (2020).

30. Jung, Y., Wen, L., Altman, A. & Ley, K. CD45 pre-exclusion from the tips of T cell microvilli prior to antigen recognition. Nat. Commun. 12, 3872, DOI: 10.1038/s41467-021-23792-8 (2021).

