## Supplementary information for "Spatial cell graph analysis reveals skin tissue organization characteristic for cutaneous T cell lymphoma"

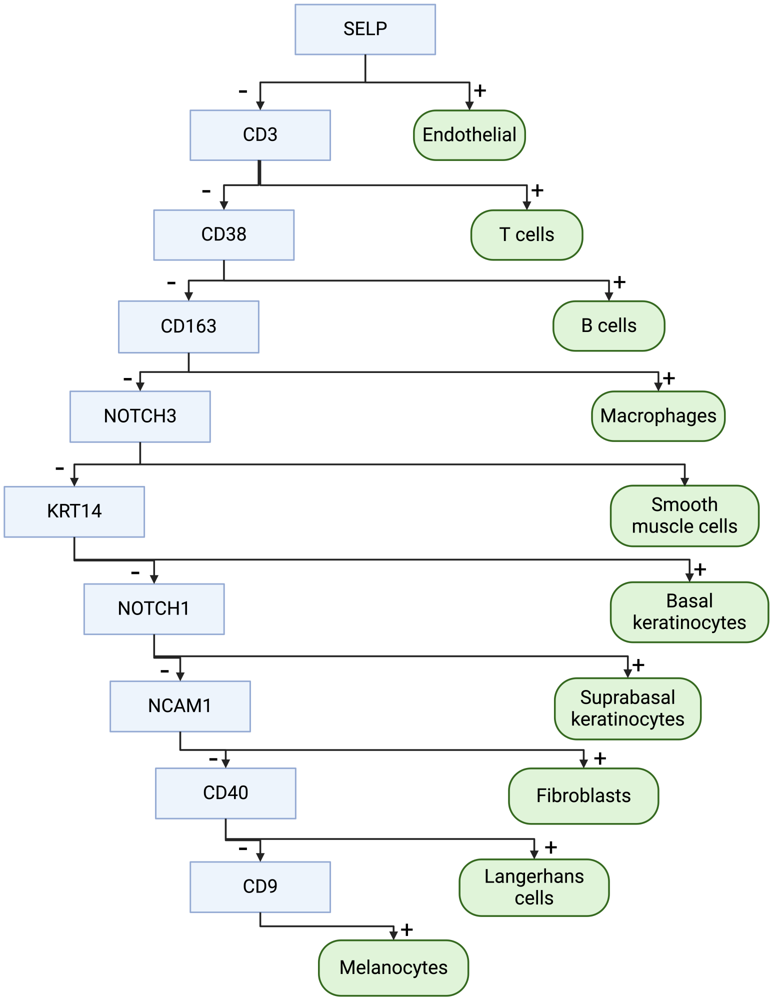


**Supplementary Figure 1.** Decision tree used for rule-based cell type assignment.

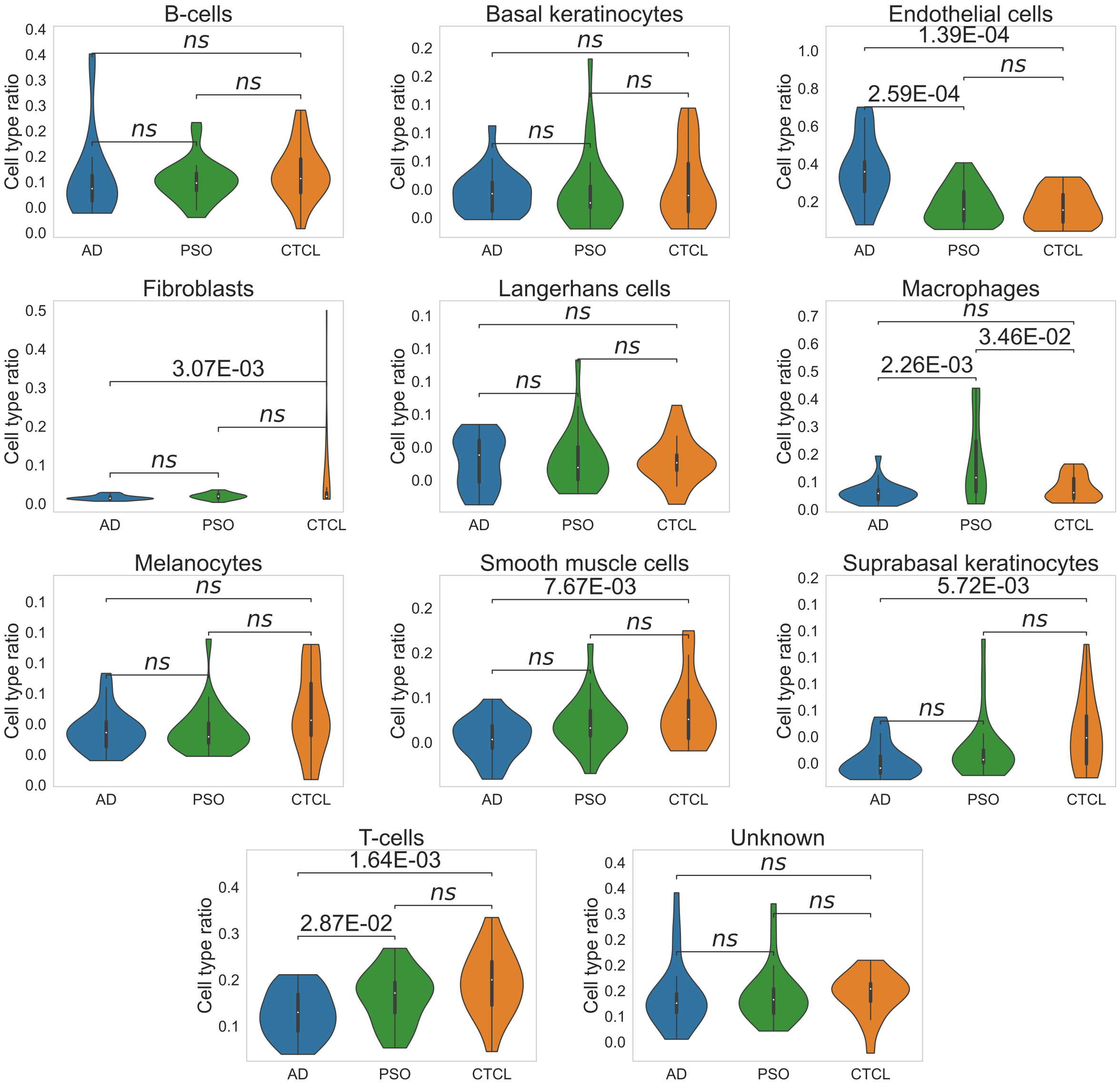


**Supplementary Figure 2.** Distributions of normalized cell type abundances across all samples, annotated with Bonferonni-corrected MWU *P*-values per condition pair.

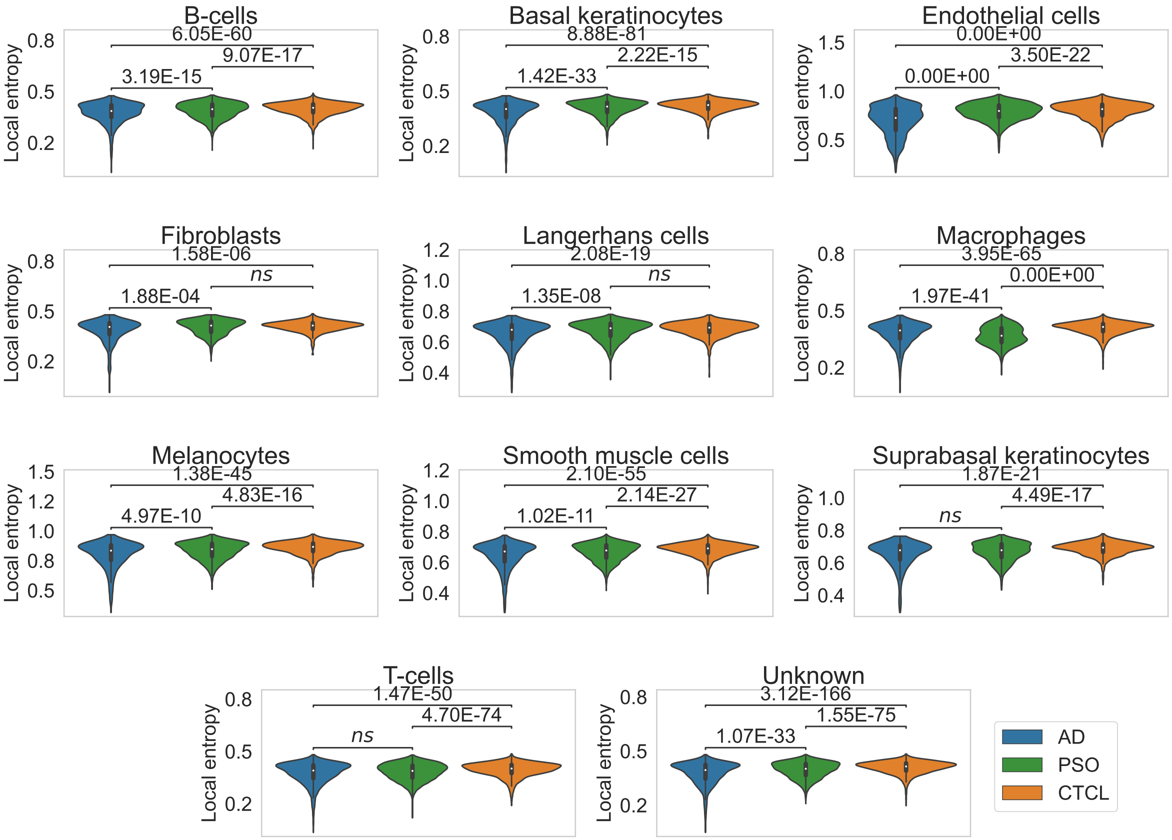


**Supplementary Figure 3.** Distributions of local entropy with radius $r = 5$, across all samples, annotated with Bonferonni-corrected MWU $P$-values per condition pair.

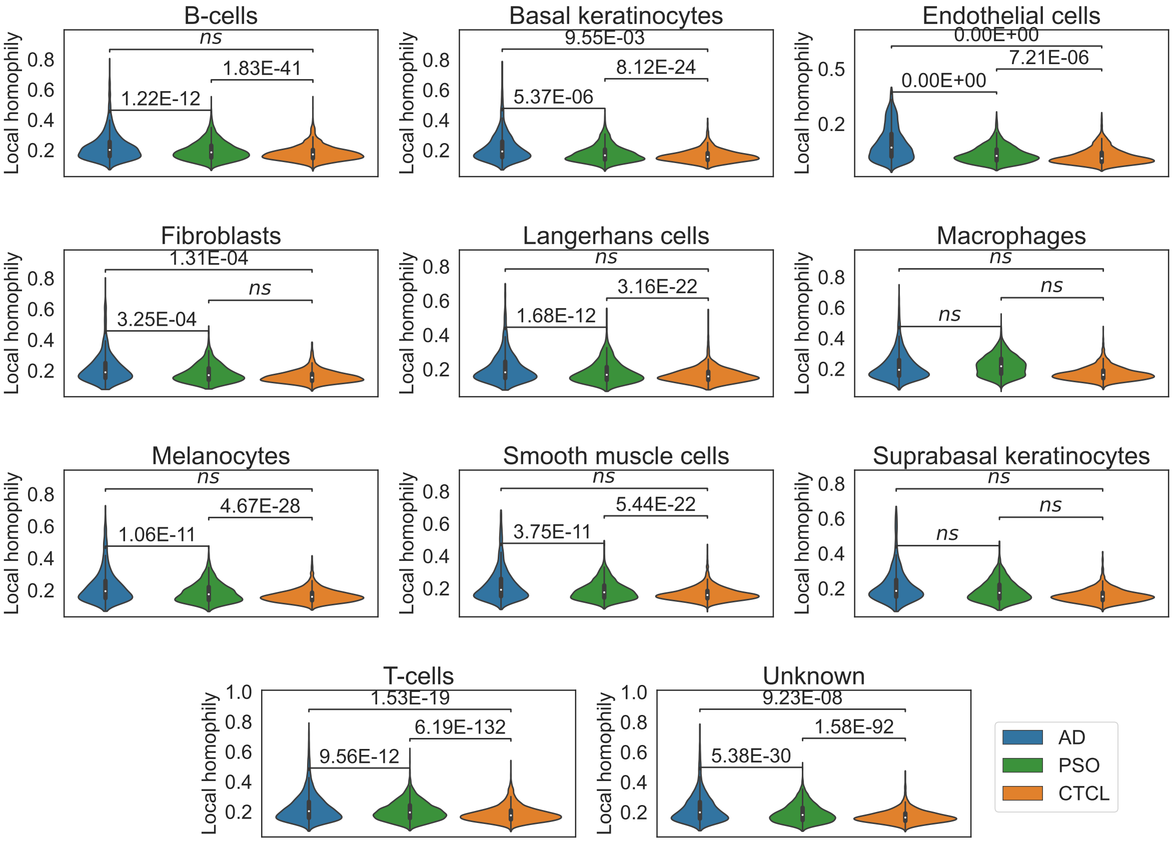


**Supplementary Figure 4.** Distributions of local homophily with radius $r = 5$, across all samples, annotated with Bonferonni-corrected MWU $P$-values per condition pair.

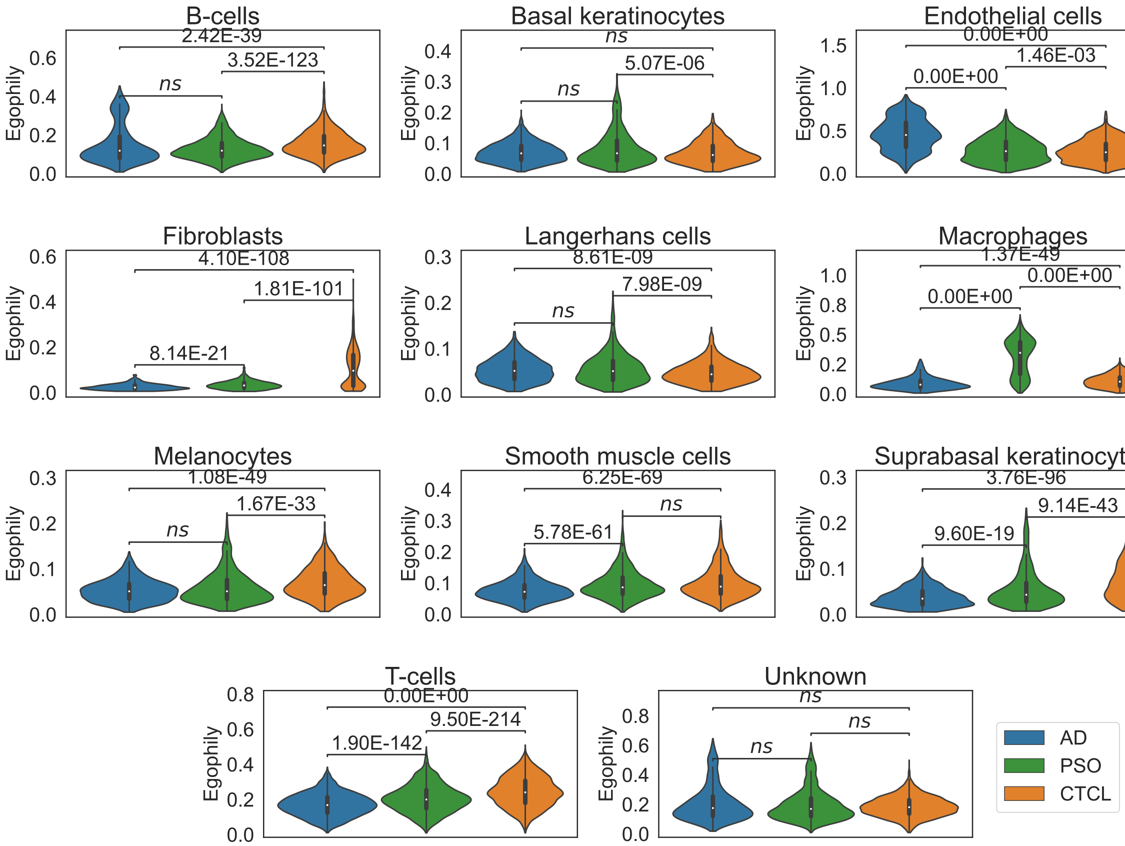


**Supplementary Figure 5.** Distributions of egophily with radius $r = 5$, across all samples, annotated with Bonferonni-corrected MWU *P*-values per condition pair.

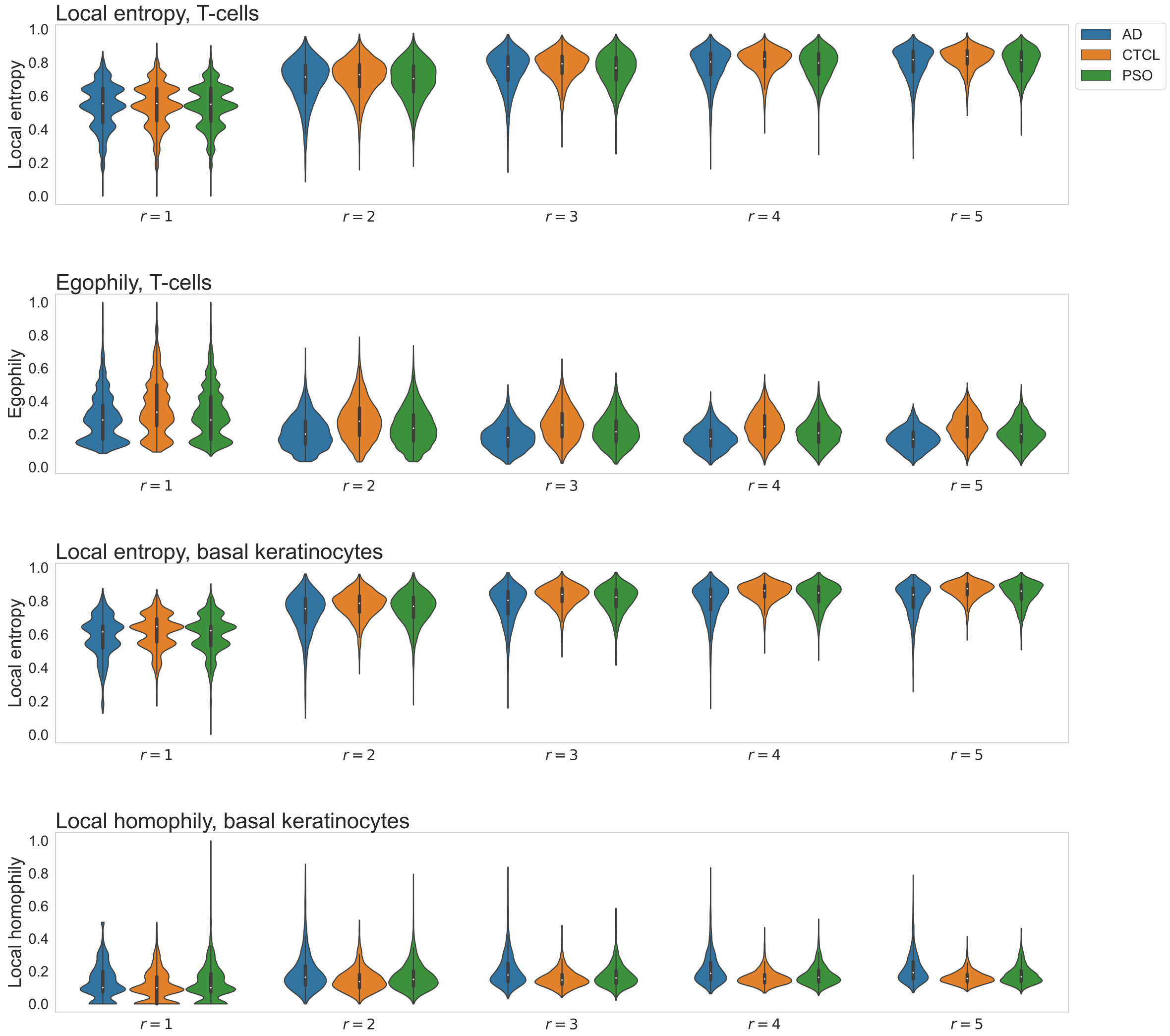


**Supplementary Figure 6.** Distributions of SHouT heterogeneity scores across radii $r \in \{1,2,3,4,5\}$ for all relevant cell types and scores as shown in Figure 3 in the main document.

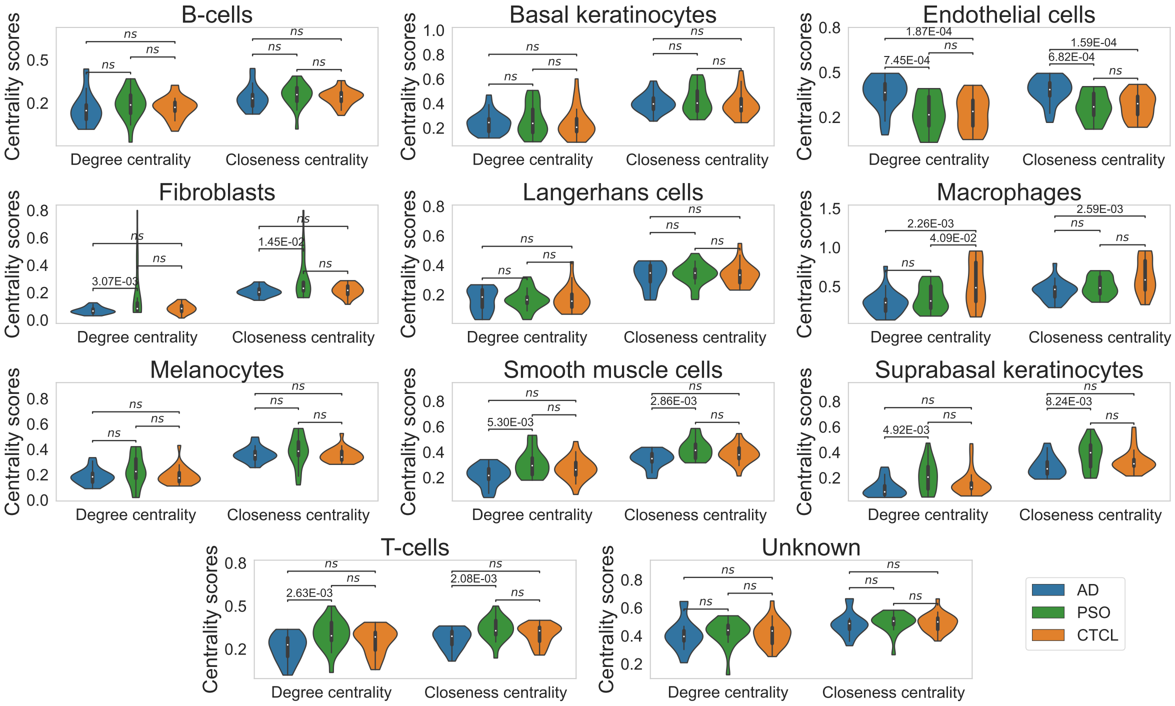


**Supplementary Figure 7.** Distributions of degree and closeness centrality scores across all samples, annotated with Bonferonni-corrected MWU *P*-values per condition pair.

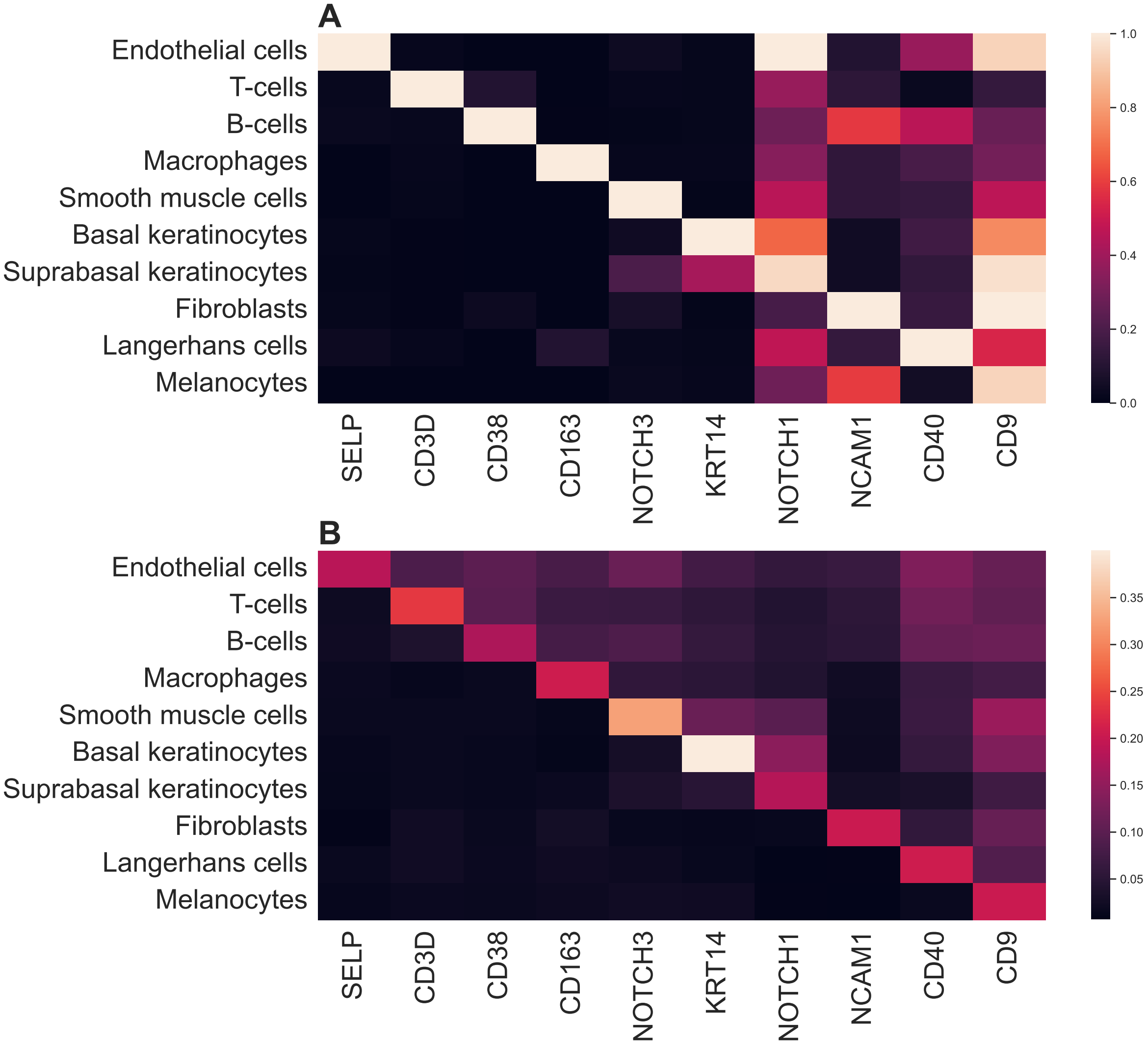


**Supplementary Figure 8.** Heatmaps showing the (A) cell type-averaged gene expression values from HPA, and (B) protein abundances from our data for the marker proteins used for cell type assignment.

**Supplementary Table 1.** Marker panel used for this study and mapping to gene names used for cell type assignment based on reference data from HPA. Since unique mapping was not possible for HLA-ABC and HLA-DR, we ran the cell type assignment approach multiple times with different combinations of possible gene name maps. Since no HLA genes were used as marker genes (see Supplementary Figure 1), all runs led to identical results.

| Original marker name | Mapped marker(s) for cell type assignment with reference data from HPA |
|---|---|
| Propodium iodide | – |
| CD11a | ITGAL |
| CD11c | ITGAX |
| CD14 | CD14 |
| CD163 | CD163 |
| CD205 | LY75 |
| CD206 | MRC1 |
| CD24 | CD24 |
| CD25 | IL2RA |
| CD29 | ITGB1 |
| CD3 | CD3 |
| CD36 | CD36 |
| CD38 | CD38 |
| CD4 | CD4 |
| CD40 | CD40 |
| CD44 | CD44 |
| CD45 | PTPRC |
| CD52 | CD52 |
| CD54 | ICAM1 |
| CD55 | CD55 |
| CD56 | NCAM1 |
| CD6 | CD6 |
| CD62P | SELP |
| CD63 | CD63 |
| CD68 | CD68 |
| CD69 | CD69 |
| CD8 | CD8 |
| CD9 | CD9 |
| CD95 | FAS |
| Cytokeratin-14 | KRT14 |
| HLA-ABC | HLA-A or HLA-B or HLA-C |
| HLA-DR | HLA-DRA or HLA-DRB1 or HLA-DRB3 or HLA-DRB4 or HLA-DRB5 |
| Notch-1 | NOTCH1 |
| Notch-3 | NOTCH3 |
| PPARgamma | PPARG |
| beta-Catenin | CTNNB1 |
